# Purification and structural characterisation of mRNA monomers, dimers, and multimers using atomic force microscopy

**DOI:** 10.64898/2026.11.20.767322

**Authors:** Alexandra L. J. Webb, Harriet Read, Thomas Minshull, Emma N. Welbourne, Prerna Bora, Lily Brewster, Manoj Pohare, Thomas E. Catley, Henrietta J. B. Smith, Preksha Hooda, Zoltán Kis, Sara Trabulo, Eivor Örnskov, Alice L. B. Pyne, Mark J. Dickman

**Affiliations:** School of Chemical, Materials and Biological Engineering, University of Sheffield, Sheffield, S1 3JD, UK; Department of Chemical Engineering, Imperial College London, Roderic Hill Building, South Kensington Campus, London SW7 2AZ, UK; Analytical Sciences, BioPharmaceuticals Development, BioPharmaceuticals R&D, AstraZeneca, Cambridge Biomedical Campus, Cambridge, CB2 0AA, United Kingdom; Advanced Drug Delivery, Pharmaceutical Sciences, BioPharmaceuticals R&D, AstraZeneca, Gothenburg, Sweden

**Keywords:** mRNA medicines, Ion-Pair Reversed-Phase HPLC, Mass photometry, Atomic force microscopy, mRNA multimers, mRNA aggregates

## Abstract

mRNA has emerged as an important new class of medicines, offering new opportunities as vaccines and therapeutics. mRNA molecules are large, highly polar and fold into complex, higher-order structures, resulting in heterogeneity. Multimerisation (aggregation) is a critical quality attribute of mRNA and may affect translational efficiency or lipid nanoparticle formulation. However, mRNA multimerisation remains poorly studied, as robust analytical methods are required for characterisation and structurally distinct populations are difficult to isolate for downstream analysis. We have developed and utilised preparative-scale, non-denaturing ion-pair reversed-phase HPLC to purify distinct mRNA monomer, dimer and multimer populations. Downstream desalting using tangential flow filtration ensured no significant perturbation of purified mRNA populations. Further characterisation of the mRNA higher-order structures was performed using Atomic Force Microscopy and subsequent automated structural analysis. mRNA dimers/multimers formed structured higher-order assemblies rather than end-to-end elongations, with mRNAs maintaining a distinct shape, even during multimerisation. Purifying distinct mRNA monomer, dimer and multimer populations establishes a platform for detailed structural, biochemical and biophysical analysis of higher-order mRNA structures. These methodologies provide insight into complex mRNA secondary, tertiary and quaternary structures. Integrating these approaches with design strategies to enhance mRNA stability and protein expression will accelerate the development of next-generation mRNA therapeutics.

## 1. Introduction

mRNA medicines offer new opportunities for potential treatments for a wide range of diseases, including vaccines against infectious diseases, cancer vaccines and protein replacement therapeutics (1–7). Analytical and purification strategies for mRNA drug substance (DS) are still being developed to enable accurate characterisation and effective separation of structurally heterogeneous mRNA populations (8–13). Higher-order mRNA species, including dimers and multimers, are critical quality attributes (CQAs) of mRNA DS (8, 9, 14) but remain relatively uncharacterised. These species may impact stability, translational efficiency, and lipid nanoparticle (LNP) formulation behaviour, highlighting the need for robust characterisation and purification methods capable of resolving structurally distinct mRNA populations.

Large-scale methods for mRNA purification include precipitation, diafiltration, and chromatographic techniques (15–26). While precipitation and membrane-based methods are widely used to concentrate RNA and remove low molecular-weight impurities, they generally lack the selectivity to remove truncated transcripts, double-stranded RNA, or structurally heterogeneous mRNA species (16, 17, 22–24). However, chromatographic purification methods offer greater selectivity, such as polythymidine (oligo-dT) affinity chromatography, size-exclusion chromatography (SEC) (25) and ion-pair reversed-phase HPLC (IP-RP HPLC) (23, 27). Oligo-dT chromatography effectively purifies full-length mRNA transcripts by binding to the poly(A) tail, thereby removing partial-length products and impurities, including residual nucleoside triphosphates (NTPs) (18, 20, 21). SEC methods also allow purification of RNAs to remove impurities, including residual NTPs, short transcripts, and DNA templates, and size-based separation for analytical characterisation of RNA (25, 26). IP-RP HPLC is often used for mRNA integrity analysis (11, 28–30). Previous studies have used IP-RP for downstream purification post-IVT to remove truncated RNA and residual pDNA template, and it is highly effective at removing dsRNA from IVT-synthesised mRNA, which can induce an innate immune response (23, 27, 31, 32).

However, methods capable of characterising and purifying discrete mRNA multimer species remain limited. SEC analysis of mRNA multimers has previously been performed, with the ultra-wide pore sizes of SEC columns enabling separation of these multimers from mRNA monomers (9, 11, 13, 33, 34) and, when coupled with multi-angle light scattering, providing molecular weight information for multimer species (9, 34, 35). However, SEC analysis typically provides lower-resolution separations, especially for mRNA >1000 nucleotides (13).

Structural characterisation of mRNA is essential for determining how structural features and higher-order structure may impact mRNA function, including translational efficiency (9, 10, 36) and encapsulation efficiency into LNPs (36). Structural analysis also provides insight into mRNA stability (9, 10), folding (10, 36, 37), and RNA interactions under different environmental conditions, such as ionic strength, cation concentration and availability, and temperature (8, 10, 38–41). Structural characterisation of native mRNA is challenging due to its large size (>1000 nts), secondary-structure heterogeneity (10, 42–44), and the distribution of multimeric conformations (8, 9). Because the intrinsic structure of RNA is conformationally flexible and dynamic (41, 43, 45), it can hinder crystallisation and particle alignment needed for high-resolution techniques, such as X-ray crystallography (37, 46). Cryogenic electron microscopy has been used to study RNA with stabilised or highly folded structures, such as ribosomes (47, 48), RNA ribozymes (49–53), viral RNA-capsid structures (54, 55), and highly folded regions of viral RNA (56, 57), although a large proportion of RNA structures studied to date are protein-RNA complexes (58). However, naked, full-length mRNA is challenging to analyse due to its long, flexible, and structurally heterogeneous nature (37, 58).

Therefore, solution-based biophysical techniques, such as small-angle X-ray scattering (SAXS) or atomic force microscopy (AFM), provide insights into mRNA conformational size and structure (36, 59–67). SAXS characterises the conformational states and molecular weight of biological macromolecules, and enables analysis under near-native conditions (63, 64). SAXS has been used to study RNA structures, and recent predictive tools can integrate SAXS data to provide insights into RNA structures and riboswitches (66, 68–72), including a 161 nt RNA riboswitch located in the 5’ UTR (70), a 94 nt aptamer, tRNAs (<100 nts) and a P4-P6 domain 158 nt RNA structural component (72). However, the large size and structural heterogeneity of mRNA limit its application, with an isolated conformational population required for accurate data (73). Predictive modelling algorithms are also being developed to determine RNA secondary structures (74, 75), with recent advances including the sequestration of Mg^2+^ within the RNA structure (73), as ions contribute to the SAXS signal and are often omitted in structural determinations. The 3D structures of long non-coding RNAs, such as braveheart (636 nts) (76), subgenomic flavivirus RNAs (∼500 nts) (77), and materially expressed gene 3 (1.6k nts), have been studied when bound to stabilising protein complexes (78).

Atomic force microscopy (AFM) provides higher spatial resolution than SAXS and performs topographical imaging of single molecules without averaging (79, 80). A key challenge for RNA imaging is the immobilisation and adsorption of the sample to the surface, which is crucial for data quality. Cations, such as Mg^2+^, Co^2+^, K^+^, and Na^+^, facilitate the association of negatively charged RNA with the mica surface, and positively charged 1-(3-aminopropyl)silatrane (APS)-coated mica is commonly used for RNA imaging (36, 59, 81). AFM has previously been used to characterise a wide range of RNA structures, such as RNase p RNA (417 nt), HIV-1 RRE RNA (350 nt) (80), metal-ion assisted folded fLuc mRNA (36) riboswitch conformers (∼250 nts) (81) 2Luc mRNA (3000 nt) (82) and A-form dsRNA (83, 84). Biophysical interactions have previously been characterised for RNA complexes, including RNA-protein (82) and RNA-lipid complexes (e.g., mRNA-LNP (85, 86)), for mRNA-DNA nanostructures and for AFM-based mRNA mapping within the cell (60).

In this study, we have developed and utilised preparative-scale, non-denaturing IP-RP HPLC for the purification of mRNA monomer, dimer, and multimers. Large quantities of pure, distinct mRNA populations suitable for further downstream studies were obtained and characterised using IP-RP HPLC and mass photometry. Further characterisation and structural studies were performed using AFM and mass photometry, with associated quantitative analysis to characterise the mRNA higher-order structures and provide important further insight into mRNA quaternary structure.

## 2. Materials and methods

### 2.1 Chemicals

Triethylammonium acetate (TEAA) (HPLC grade, Glen Research), acetonitrile (ACN) (HPLC grade, Fisher), water (HPLC grade, Fisher), and a 2 M magnesium chloride solution (Sigma), 10x PBS solution (Fisher) were used. *In vitro* transcription (IVT) used ATP, CTP, GTP, UTP (Roche), inorganic pyrophosphatase (IPP, Roche), T7 RNA polymerase (Roche), RNase inhibitor (Roche), DNase I (Roche), HEPES (Gibco), NaCl (Invitrogen), magnesium acetate (Thermo Scientific), DTT (Thermo-Fisher), spermidine (Sigma-Aldrich), Triton X-100 (Merck), DNA template (GenScript), nuclease-free water (Ambion). Oligo-dT purification of mRNA used 1 M sodium phosphate (pH 7.2), 0.5 M EDTA (Invitrogen), 8 M guanidinium hydrochloride (GuHCl, Thermo Scientific), 1 M NaOH (Scientific Laboratory Supplies), and CIMmultus^TM^ Oligo dT18 4 mL monolithic column, C12 linker with 2 µm channels (Sartorius). High-range Riboruler RNA ladder (Thermo Fisher) was used for sizing. Tangential flow filtration (TFF) purification used HEPES (1 M, Fisher Scientific), Tris-HCl (1 M, Invitrogen), and NaOH (1 M, Scientific Laboratory Supplies). Aminopropylsilatrane (APS, Sigma-Aldrich)

### 2.2 In vitro transcription and oligo-dT purification of the mRNA sample

IVT components (NTPs, 10x buffer, IPP, RNase inhibitor, DNA template, T7 polymerase, and nuclease free water) were incubated at 37 °C for 2 hours. Samples were diluted to 0.4 mg/ml with binding buffer (50 mM sodium phosphate (pH 7.2), 400 mM GuHCl, and 5 mM EDTA). Oligo-dT purification was performed using an ÄKTA Pure 25 system (Cytiva) with oligo dT18 (C12)-4 monolith (CIMmultus). Purified RNA was eluted with nuclease free water, collected in 1 mL fractions into a 96-well plate, and RNA peak fractions were pooled at the end. Quantification was performed using a NanoDrop^TM^ One UV-vis spectrophotometer (Thermo Fisher Scientific) with absorbance at 260 nm.

### 2.3 Preparative purification of mRNA monomer, dimer, and multimer species using non-denaturing IP-RP chromatography

For all preparative IP-RP chromatography, an ÄKTA avant system (Cytiva), a DNAPac RP semi-preparative column (4 μm, 21.2 × 100 mm) (Thermo Fisher Scientific), a mobile-phase Caloratherm preheater (Selerity Technologies) and a 2 mL sample loop were used. Flow rate was 6 mL/min, and UV detection at 260 nm. 2 mL fractions were eluted according to peak-detection parameters and collected into a 96-well plate. UNICORN (Cytiva) software was used for method editing, system control, and data acquisition.

#### Gradient 1 (denaturing)

Mobile phase A: 0.1 M TEAA and B: 0.1 M TEAA, 25% ACN. Column oven temperature was 70 °C. The gradient consisted of a 15-minute wash step at 30% B, a linear elution step from 30% to 70% over 35 minutes, a 15-minute wash step at 100% B, and a 15-minute column equilibration at 30% B.

#### Gradient 2 (non-denaturing)

To separate mRNA monomer, dimer, and multimer species. Mobile phase A: 0.1 M TEAA, 1 mM Mg^2,+^ and mobile phase B: 0.1 M TEAA, 1 mM Mg^2+^, 25% ACN. Column oven temperature was 50 °C. The gradient consisted of a 15-minute wash step at 30% B, a linear elution step from 30% to 60% over 35 minutes, a 15-minute wash step at 100% B, and a 15-minute column equilibration at 30% B.

### 2.4 Tangential flow filtration

System 1 - A KrosFlo® Repligen 2i (KR2i) TFF system was used with 300 kDa Sartocon Slice 50 Hydrosart (cellulose) cassettes (Sartorius). The column was washed twice with Type I water (flushing volume of 2.0 mL/cm^2^ of membrane area) and sample buffer. The feed tank was then filled with the sample, and the accumulated masses of the retentate and permeate were recorded. The software was KF Comm 2 (Repligen). System cleaned with 0.1 M NaOH and deionised water, and mass calibration was performed.

System 2 - μPulse® system (Formulatrix) was used with 100 kDa regenerated cellulose (RC) filter chips (Formulatrix). Buffer exchange enabled automatic diafiltration for buffer exchange/desalting. System was cleaned with 0.1 M NaOH and deionised water, and mass calibration was performed.

### 2.5 Analytical IP-RP HPLC analysis

IP-RP HPLC analysis was performed on U3000 and Vanquish HPLC systems (Thermo Fisher) using a ProSwift RP 3U 4.6 × 40 mm column (Thermo Fisher). Mobile phase A was 0.1 M TEAA, 1 mM Mg^2+^, and mobile phase B was 0.1 M TEAA, 1mM Mg^2+^, 25% ACN. The gradient consisted of a linear step from 20-25% over 1 minute, a non-linear extension to 70% B over 14 minutes, and a 5-minute wash step at 100% B. Flow rate was 0.50 mL/min, and UV detection was at 260 nm.

For non-linear gradient:

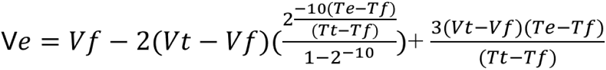

### 2.6 Mass photometry analysis

A TwoMP mass photometer (Refeyn) was used. Glass coverslips (24 × 50 mm, Epredia, UK) were rinsed twice with Milli-Q water, methanol, and isopropanol, then dried under nitrogen. Coverslips were coated with 0.01% poly-L-lysine for 5 minutes, then dried under nitrogen. RNA samples were diluted in 1X PBS to <5 ng/µL. Droplet dilution measurements were performed using 1 X PBS. The Riboruler High Range RNA Ladder served as the calibration standard for mRNA sizing. Measurements were recorded for 60 seconds. Data acquisition and analysis were performed using Acquire MP and Discover MP software (Refeyn), respectively. The average multimer size from the distribution was calculated by:

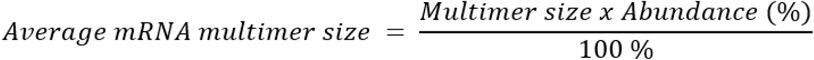

### 2.7 AFM Imaging and automated processing for analysis

All AFM measurements were carried out using a commercial AFM (Dimension XR FastScan, Bruker) in PeakForce QNM mode. Measurements were performed in liquid at room temperature. Standard Fastscan-D probes (Bruker) were used with nominal values k ≈ 0.25 N m^-1^ and f0 ≈ 110 kHz at a PeakForce tapping frequency of 8 kHz. Typical forces applied by the AFM tip to the sample ranged between 50 - 300 pN; these forces were kept low to avoid any sample damage. To obtain suitable AFM substrates for RNA immobilisation, mica discs (diameter 0.5 cm^2^) were functionalised with APS (Sigma Aldrich), as described by Shlyakhtenko et al. (59). For all RNA samples, 1-2 μL of ∼10 ng/uL RNA was deposited onto the APS-mica surface in 20 μL of 150 mM KCl, 20 mM HEPES (pH 7.4) buffer. The RNA was incubated for 5 minutes before being rinsed with the final imaging buffer, 50 mM NaCl, 20 mM HEPES (pH 7.4). The samples were immediately transferred for AFM imaging in 20 uL of imaging buffer to ensure they remained hydrated. Post-processing and analysis of the AFM images were performed using the open-source software Topostats (87). Height thresholding was performed to segment individual molecules as binary masks. Each molecule was then assigned a grain index, and descriptors corresponding to the shape and size of the molecule were calculated. These included the volume, smallest bounding area (SBA, the area of the smallest possible bounding box around the molecule, which is not constrained to the Cartesian axes), minimum and maximum feret diameter (minimum/maximum distance between two parallel planes restricting the molecule), aspect ratio (ratio of min: max feret diameter), and mean height of the mRNA molecules.

Normality tests were performed on the AFM data for both eGFP and fLuc mRNA using a Shapiro-Wilk normality test, where p > 0.05 indicated no significant deviation from normality, to determine which statistical test was appropriate. The normally distributed data were analysed using one-way ANOVA followed by Tukey’s multiple-comparison test. For eGFP, the number of mRNA molecules analysed in purified samples was: monomer n=100, dimer n=104 and multimer n=87. For fLuc, the number of mRNA molecules analysed in purified samples was: monomer n=101, dimer n=111 and multimer n=102. Data that showed significant deviations from normality (p < 0.05) were analysed using Kruskal-Wallis significance tests, followed by Dunn’s multiple comparison test. AFM measurements are reported as mean ± standard deviation (n < 500 molecules). Significance was assigned as such, p < 0.0001 ****, p < 0.001 ***, p < 0.01 **, p < 0.05 * and ns= not significant (p > 0.05).

## 3. Results and Discussion

### 3.1 Preparative-scale, non-denaturing ion-pair reversed-phase chromatographic purification of mRNA monomer, dimer, and multimers

We have previously developed an analytical scale, non-denaturing IP-RP HPLC method for the separation and characterisation of mRNA multimers (aggregates) (8). Incorporation of 1 mM Mg²⁺ into the 0.1 M TEAA mobile phase enabled stabilisation and resolution of monomeric, dimeric, and multimeric species across multiple mRNAs of varying lengths and sequences (8). To obtain sufficient quantities of purified mRNA monomers, dimers, and multimers for downstream studies, it was necessary to scale up the non-denaturing IP-RP HPLC and implement downstream desalting. Here, we developed a preparative, non-denaturing IP-RP chromatographic method using an ÄKTA avant chromatography system and a semi-preparative macroporous PS-DVB stationary-phase column (DNAPac RP, 4 μm, 21.2 × 100 mm). Precise temperature control of the column was essential to facilitate both mRNA denaturation at 70 °C and multimer formation at lower temperatures. This is challenging due to the high flow rates (6 mL/min) and large column volumes involved. Therefore, the IP-RP separations were performed using both a column oven and a dedicated mobile-phase pre-heater suitable for the high flow rates required in preparative-scale chromatography. The analysis and purification of eGFP mRNA (4.2 mg) under denaturing conditions at 70 °C using 0.1 M TEAA on the preparative scale is shown in Figure 1A.

**Figure 1.**
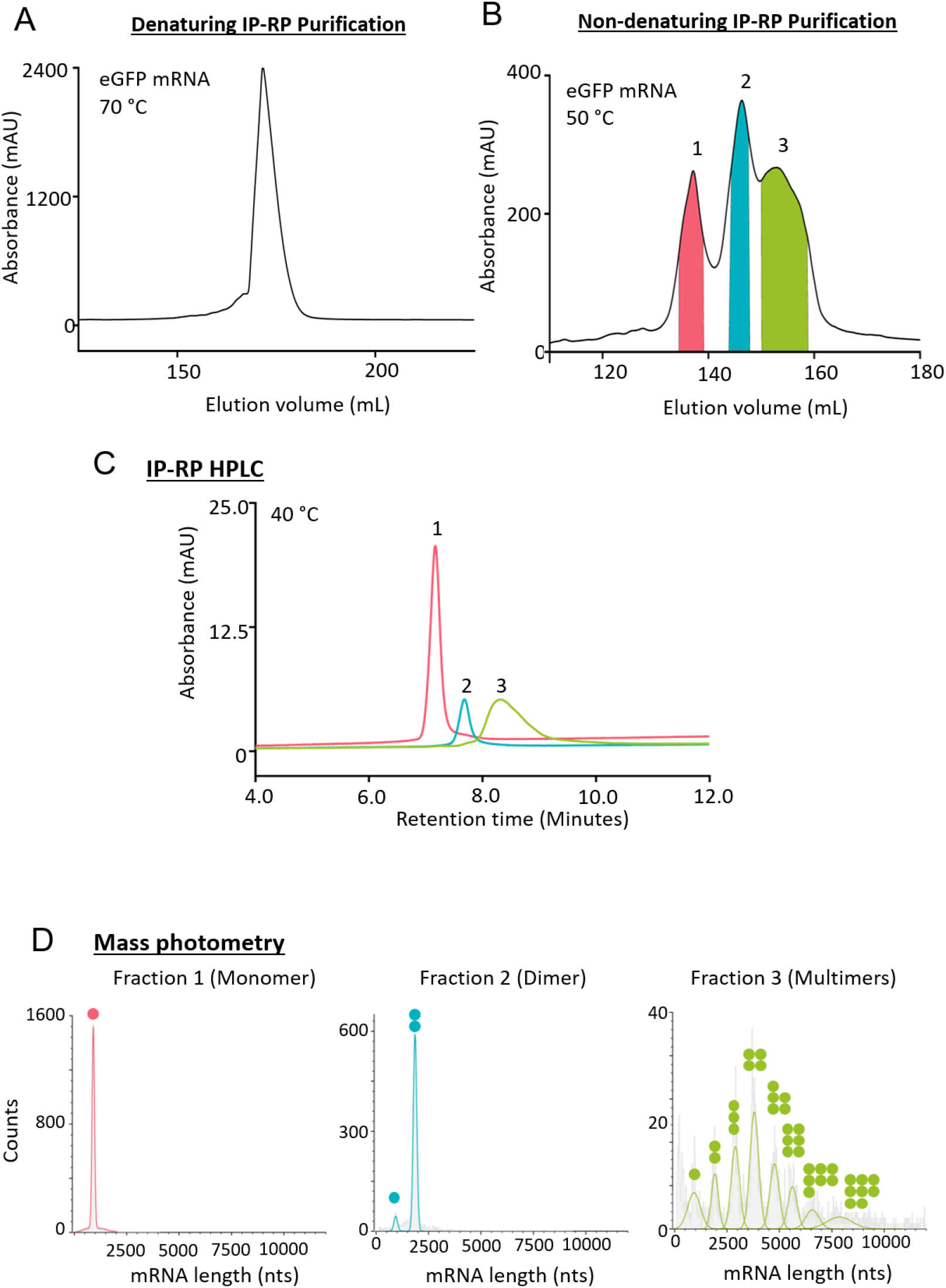
A) Preparative denaturing IP-RP purification of eGFP mRNA. 4.2 mg of eGFP mRNA was analysed using a DNAPac RP (21.2 × 100 mm) column at 70 °C, UV detection 260 nm using preparative gradient 1. B) Preparative non-denaturing IP-RP purification of eGFP mRNA. 4.2 mg of eGFP was analysed at 50 °C using a DNAPac RP (21.2 × 100 mm) column using preparative gradient 2. Fractions (1–3) were collected, corresponding to purified eGFP monomer, dimer and multimer populations. C) Non-denaturing IP-RP HPLC analysis of fractions 1, 2 and 3 shows singular populations at alternative retention times corresponding to monomer, dimer and multimer populations. D) MP analysis of purified fractions 1, 2 and 3. The corresponding sizes and relative abundance show fraction 1 is pure monomer species, fraction 2 is the majority dimer species, and fraction 3 is a mix of multimer species. Alt text: Four figure panel showing purification and characterisation of eGFP mRNA. A) Preparative scale denaturing IP-RP purification of eGFP mRNA, showing single peak at approximately 170 mL elution volume. Absorbance at 260 nm is plotted against elution volume. B) Preparative non-denaturing IP-RP purification of eGFP mRNA with three peaks collected as fractions, corresponding to monomers, dimers and multimers. C) Non-denaturing IP-RP HPLC analysis of fractions 1, 2 and 3 shows singular populations at alternative retention times corresponding to monomer, dimer and multimer populations. D) MP analysis showing monomer fraction is 100% monomeric, dimer fraction >90% dimeric and multimer sample has distribution of species from monomer to nonamer.

We then exchanged the mobile phase to 0.1 M TEAA, 1 mM Mg^2+^, with the column and mobile phase pre-heater at 50 °C to avoid disrupting RNA: RNA interactions, as mRNA multimerisation is dependent on lower temperatures (<60 °C). We had previously observed at the analytical scale that using column oven temperatures of 50-55 °C increases dimer/multimer formation in eGFP mRNA, suggesting that partial unfolding of the mRNA secondary structure promotes dimer formation (8). Preparative scale, non-denaturing IP-RP separation (50 °C) of 4.2 mg eGFP mRNA generated three distinct chromatographic peaks, which were collected in 2 mL fractions and pooled into fractions 1, 2 and 3 (Figure 1B). Further characterisation of the fractions was performed using non-denaturing IP-RP HPLC (Figure 1C). Fractions 1, 2, and 3 had increasing retention times, corresponding to the mRNA monomer, dimer, and multimer populations (8). Further characterisation was performed using mass photometry to enable sizing and relative abundance of the different mRNA higher-order structure populations (Figure 1D and Supplementary Figures S1 and S2). The mass photometry results showed that fraction 1 contained 100% mRNA monomer, fraction 2 contained >90% mRNA dimer, and fraction 3 contained a mixed population of mRNA higher-order multimers, up to and including a 9-mer (Figure 1D and Supplementary Table S1). Overall, the method enables effective separation and purification of mRNA monomer, dimer and higher-order multimer populations suitable for downstream structural and functional studies.

### 3.2 Desalting of mRNA multimers using tangential flow filtration (TFF)

The purified mRNA fractions using IP-RP chromatography were eluted in 0.1 M TEAA, 1 mM Mg²⁺ with ∼ 12.5% ACN, and therefore required further downstream processing to remove residual ion-pairing reagent, ACN, and free Mg²⁺. This was necessary to minimise accelerated hydrolytic degradation of mRNA, which is catalysed by Mg²⁺ ions (38, 88, 89). The removal of the ion-pairing reagent and organic solvent is also required for any downstream studies, including LNP formulation and cell transfections. Moreover, any downstream desalting methods must not dissociate or perturb the purified mRNA monomer, dimer, and multimer populations to enable further downstream studies and comparisons of the mRNA higher-order structures. Two alternative TFF (diafiltration) methods were compared: the μPulse system using 100 kDa RC filter chips and the KR2i system using 300 kDa Sartocon Slice 50 Hydrosart cassettes. Diafiltration was performed on IP-RP-purified eGFP mRNA monomer, dimer and multimer fractions. Analysis using non-denaturing IP-RP HPLC and mass photometry (MP) of samples before and after diafiltration was performed to determine any changes in mRNA populations.

Diafiltration on the KR2i system did not significantly alter the relative abundance of mRNA monomer, dimer and multimers in each of the fractions analysed. The non-denaturing IP-RP HPLC analysis (Figure 2A) shows that fractions 1, 2, and 3 purified with the KR2i system remain as single discrete mRNA populations, with retention times consistent with those of the pre-TFF monomer, dimer and multimer species, respectively.

**Figure 2.**
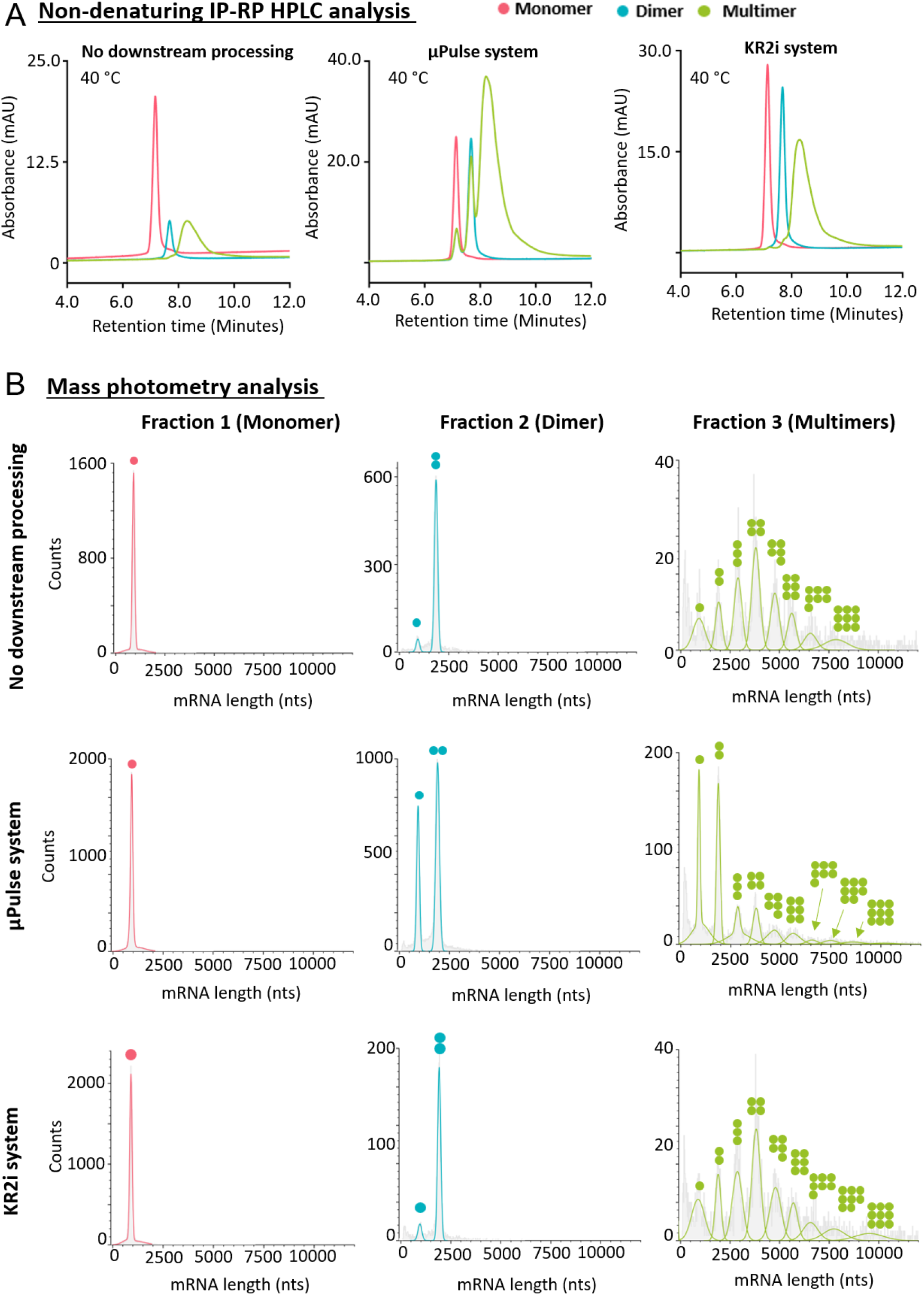
A) Non-denaturing IP-RP HPLC analysis and B) MP analysis of eGFP fractions 1, 2 and 3 before downstream processing and post-TFF with KR2i and μPulse systems. Alt text: Two panels showing comparison of TFF systems using A) IP-RP HPLC and B) MP. Both panels indicate μPulse system perturbs dimer and multimer structures, while KR2i does not.

However, diafiltration using the μPulse system altered the relative abundance of mRNA populations within the higher-order fractions (2, 3). In fraction 2 (purified mRNA dimer), an increase in the relative abundance of the mRNA monomer was observed by both IP-RP HPLC analysis (Figure 3A) and MP (Figure 2B). Similarly, fraction 3 post-diafiltration showed the presence of, and an increased relative abundance of, mRNA monomers and dimers (Figure 2). Therefore, under these conditions, using this diafiltration system, a proportion of mRNA dimers and multimers dissociated into monomers. The KR2i system was therefore used for all future experiments. Following TFF purification using the KR2i system, samples were stored at -80 ℃, and short-term stability studies showed that the mRNA monomer, dimer, and multimer fractions were not significantly altered during short-term storage.

**Figure 3.**
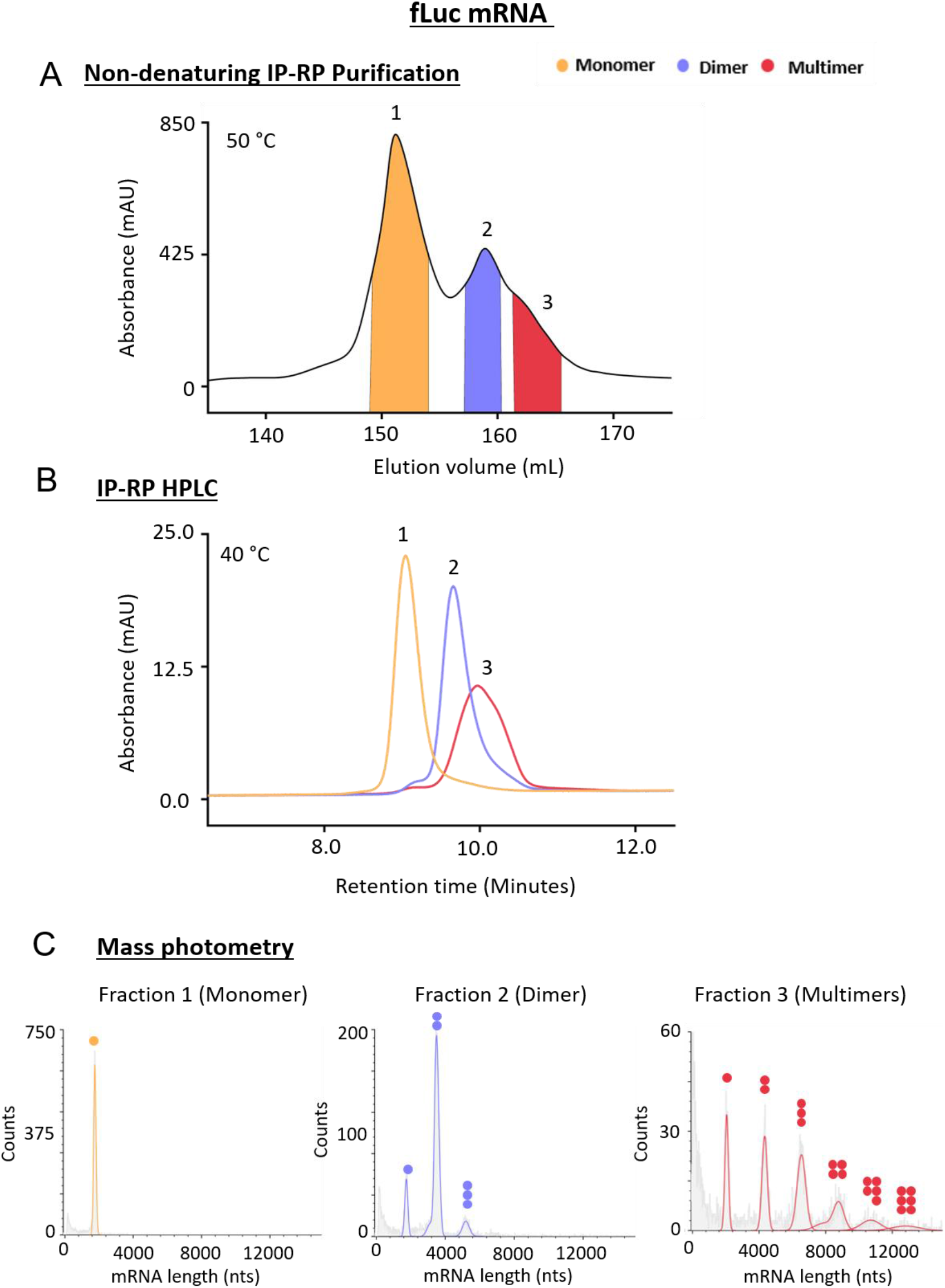
A) Preparative non-denaturing IP-RP purification of fLuc mRNA. 3.2 mg of fLuc was analysed at 50 °C using a DNAPac RP (21 × 100 mm) column using preparative system 2. Fractions (1–3) were collected, corresponding to purified fLuc monomer, dimer and multimer populations. B) Non-denaturing IP-RP HPLC analysis of fractions 1, 2 and 3 shows singular populations at alternative retention times corresponding to monomer, dimer and multimer populations. C) MP analysis of purified fractions 1, 2 and 3. The corresponding sizes and relative abundance show fraction 1 is pure monomer species, fraction 2 is the majority dimer species, and fraction 3 is a mix of multimer species. Alt text: Three figure panel showing purification and characterisation of fLuc mRNA populations. A) Preparative non-denaturing IP-RP purification of fLuc mRNA with three peaks collected as fractions, corresponding to monomers, dimers and multimers. B) Non-denaturing IP-RP HPLC analysis of fractions 1, 2 and 3 shows singular populations at alternative retention times corresponding to monomer, dimer and multimer populations. C) MP analysis showing monomer fraction is 100% monomeric, dimer fraction >80% dimeric and multimer sample has distribution of species from monomer to hexamer.

Following successful preparative-scale non-denaturing IP-RP purification of eGFP mRNA monomers, dimers, and multimers, combined with TFF purification of the corresponding mRNA populations, further work was performed to analyse and purify an alternative mRNA (fLuc mRNA). Following non-denaturing IP-RP chromatography of 3.2 mg of fLuc mRNA (see Figure 3A), the corresponding fractions 1, 2, and 3 were collected, diafiltrated and subsequently analysed using IP-RP HPLC and MP (Figures 3B and C and Supplementary Figure S3). The population distribution showed that for fLuc mRNA under the conditions analysed, the mRNA monomer species (fraction 1) was the most abundant conformer under these conditions. Furthermore, compared to eGFP mRNA, the relative abundance of the fLuc mRNA multimers is lower, indicating RNA: RNA interactions are less favourable under these conditions, and that mRNA self-association is weaker.

### 3.3 AFM imaging of purified mRNA monomer, dimer and multimers

Following successful purification and desalting of the mRNA monomer, dimer, and multimer populations, further structural characterisation was performed using AFM. mRNA molecules were deposited onto APS-mica discs for imaging in buffer solution (see Supplementary Figure S4). APS mica was used to minimise surface charge induced structural changes and to enable salt concentrations to be controlled. Applied forces to the sample by the tip were kept low (∼pN) to ensure tip-induced deformation or sample damage was minimised.

To further investigate the mRNA higher-order structures purified using IP-RP chromatography, AFM images of both eGFP and fLuc mRNA samples were analysed (see Figures 4 and 5 and Supplementary Figure S5). Comparison of the AFM images (Figures 4A and 5A) revealed differences in molecular organisation. Image analysis revealed distinct, statistically significant structural differences between the mRNA monomer, dimer, and multimer samples, most notably in the overall shape and size, area, volume and length of the molecules (see Figures 4B and 5B and Supplementary Tables S2 and S3). Numerical results of statistical testing are reported in Supplementary Tables S4-S7. Both eGFP and fLuc mRNA monomer samples consisted of individual mRNA molecules, whereas the dimer samples predominantly consisted of two associated mRNA molecules which form a larger higher-order structure. The multimer populations contained higher-order assemblies formed of multiple associated mRNA molecules. Furthermore, AFM imaging of the multimer samples revealed a mixed population of species with a broad size distribution, aligning with the range of mRNA species detected by MP (Figure 3C).

**Figure 4.**
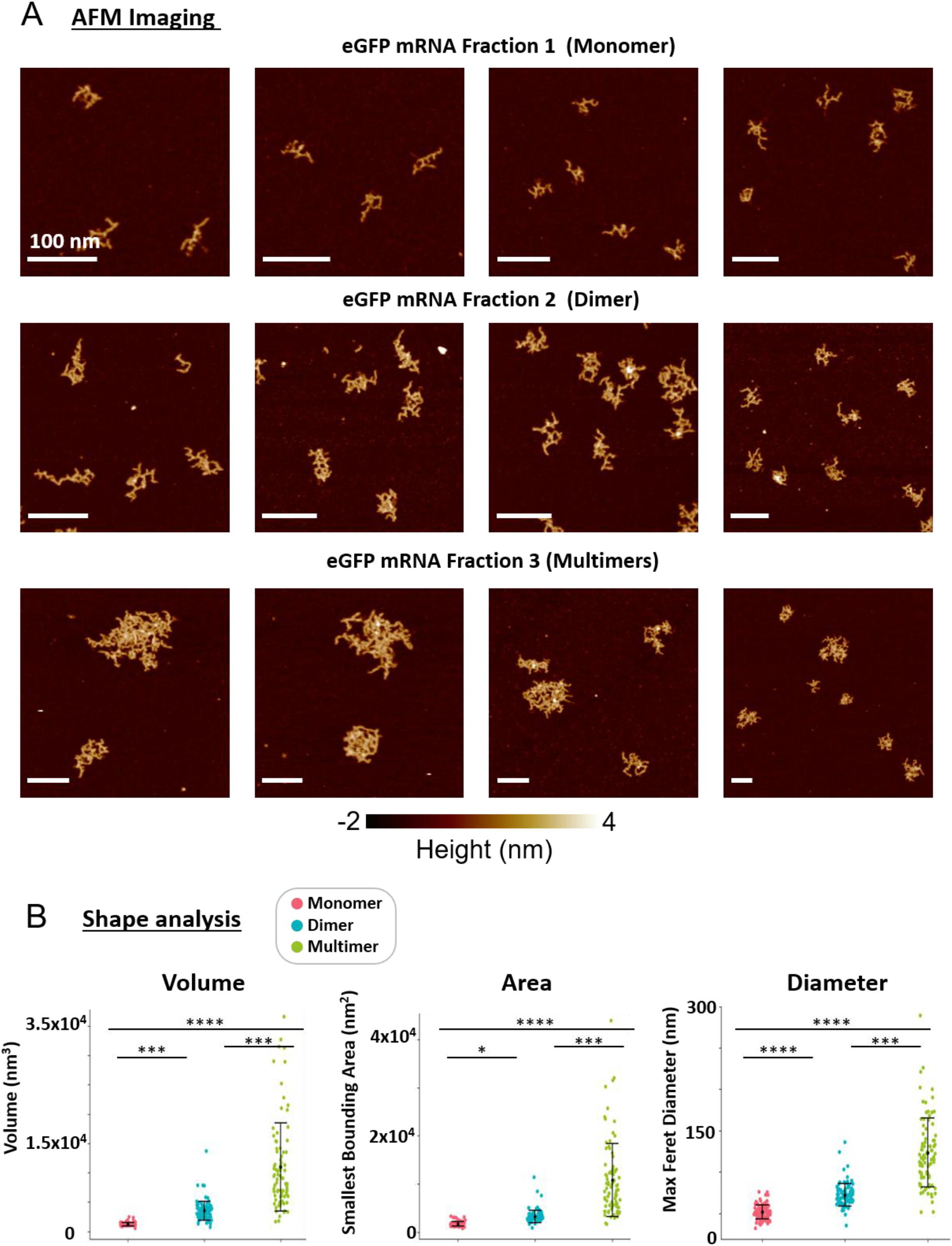
A) AFM imaging of purified eGFP monomer, dimer and multimer populations. B) Topographical analysis of volume (nm^3^), smallest bounding area (SBA, nm^2^) and maximum feret diameter (nm) for purified eGFP monomer, dimer and multimer species. Alt text: Two panels showing AFM images and shape analysis of eGFP mRNA monomers, dimers and multimers. A) Representation AFM images of monomers, dimers and multimers showing increasing complex mRNA assemblies. B) Scatter plots comparing volume, SBA and maximum feret diameter of purified monomer, dimer and multimer populations. All three measurements increase with assembly size, with statistical significance indicated above the plots (reported in supplementary information).

**Figure 5.**
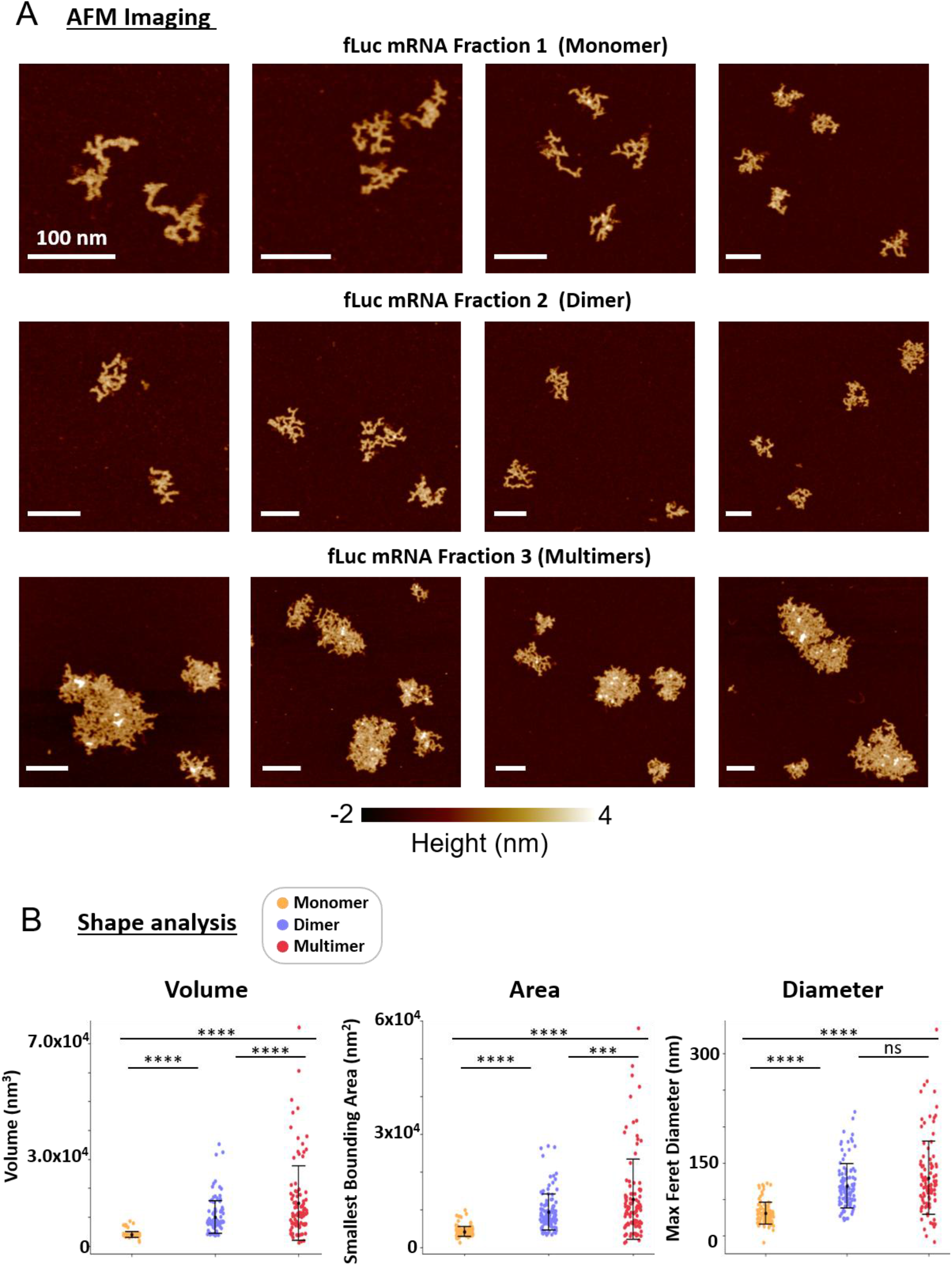
A) AFM imaging of purified fLuc monomer, dimer and multimer populations. B) Topographical analysis of volume (nm^3^), smallest bounding area (SBA, nm^2^) and maximum feret diameter (nm) for purified fLuc monomer, dimer and multimer species. Alt text: Two panels showing AFM images and shape analysis of fLuc mRNA monomers, dimers and multimers. A) Representation AFM images of monomers, dimers and multimers showing increasing complex mRNA assemblies. B) Scatter plots comparing volume, SBA and maximum feret diameter of purified monomer, dimer and multimer populations. Volume and SBA increase across the three populations, while maximum Feret diameter shows overlapping distributions between dimers and multimers with statistical significance indicated above the plots (reported in supplementary information).

### 3.4 Comparative structural analysis of mRNA higher-order structures using AFM

#### 3.4.1 mRNA dimer analysis

The IP-RP purification method above shows clearly separated mRNA populations. To gain insights into the conformational makeup of these distinct populations, and to obtain further structural information on the mRNA samples, we performed in-depth image analysis of AFM topographic images. This analysis revealed significant differences between the monomer, dimer, and multimer samples. Shape and size analysis of individual mRNA molecules was performed, extracting the volume (nm^3^), smallest bounding area (SBA) (nm^2^), maximum feret diameter (nm), mean height (nm) and aspect ratio (min: max feret diameter ratio) via TopoStats (87). For both eGFP and fLuc mRNA, the mean averaged dimer and multimer sample measurements were compared to the mean averaged monomer sample, as the species are aggregates of the monomer unit (see Figure 4 and 5).

The eGFP dimer species has a larger volume (2.9 times), smallest bounding area (1.9 times), maximum feret diameter (1.4 times), and height (1.42 times) than the mRNA monomer species (see Figure 4). The doubling of the mean SBA of the dimer species is consistent with the immobilised footprint of 2 associated monomer units. However, we observe that the dimer structure does not just adopt a simple end-to-end elongation. The increase in maximum feret diameter indicates a moderate extension of the longest molecular dimension and a moderate increase in height, albeit statistically significant from the monomer, indicating some level of overlapping or stacking of structure (see Supplementary Figure S6 and Supplementary Table S5). In comparison, the volume of the dimer is more than double that of the monomer. It is proposed that the increased volume reflects a more open secondary-structure conformation of the monomer units associated in the dimer assembly. The model of RNA adopting a more open conformation upon dimerisation or multimerisation has been previously observed for HIV-1 genomic RNA (90–95), bacteriophage ø29 pRNA (96–100), and bicoid mRNA (101, 102). In addition, we have also previously reported temperature-dependent eGFP mRNA dimer assembly, indicating that conformational changes promote mRNA dimerisation. Notably, the aspect ratios of the eGFP mRNA monomer and dimer are very similar, 0.735 and 0.732, respectively, resulting in a non-statistically significant difference, indicating that the monomer units within the dimer preserve the same lateral proportions as the monomer and do not become more elongated, which would be expected with end-to-end dimer formation.

The fLuc mRNA dimer species exhibits a larger volume, SBA, and maximum feret diameter than the monomer species by 2.5 times, 2.2 times, and 1.5 times, respectively (Figure 5B), with parameters exhibiting a highly significant difference (see Supplementary Table S7). SBA analysis is consistent with the eGFP mRNA, with the immobilised footprint approximately doubling for the dimer. As with eGFP mRNA dimer, the fLuc mRNA dimer volume is more than double that of the respective monomer, indicating that the two monomer units adopt a more open secondary structure conformation for dimer formation. The maximum feret diameter is less than double but still larger than that of the purified monomer mRNA. The mean height of the dimer increases by only 1.1 times that of the monomer, indicating minimal overlapping or stacking. The aspect ratios of the monomer and the dimer are both 0.723, indicating no significant difference in overall shape. Again, we note that the dimer does not form an end-to-end elongation, rather it forms a more compact structure, similar to the eGFP mRNA.

Overall, we see that mRNA dimers maintain the same moderately elliptical shape as monomers and do not extend in a single plane. These data are consistent with compact, ordered assemblies mediated by specific intermolecular interactions rather than nonspecific interactions. eGFP forms a dimer with a greater relative increase in volume and height upon dimerisation than fLuc, indicating a more open conformation and more partial stacking occurring from monomer to dimer structure for eGFP mRNA. The differences in eGFP and fLuc dimer formation are likely due to the nature of the folded secondary structure of the mRNAs having an open structure, which impacts the availability of nucleotide bases for intermolecular bonding.

#### 3.4.2 mRNA multimer analysis

Further work was carried out to investigate the structure and formation of mRNA multimers. MP analysis of the eGFP multimer sample showed that the size distribution of mRNA species ranged from monomer to nonamer (9-mer), but corresponded to an average of ∼4.23 monomer units, approximately a tetramer (4-mer) (Figure 1D). AFM imaging of eGFP and fLuc mRNA multimers yielded heterogeneous populations with a broad size distribution, consistent with those detected using MP. The eGFP multimer sample had an average volume, SBA and max feret diameter ∼9.0, ∼6.4 and ∼2.4 times larger than the monomer, respectively. The multimer also had an average height ∼1.4 times higher than the monomer. As with the dimer species, multimers adopt laterally associated assemblies rather than extending in one direction. The structures increase in size, but maintain a tight conformation which extends across the surface. The aspect ratio of the average eGFP multimers is 0.753, very similar to the monomer (0.735), with no statistically significant difference (see Supplementary Tables S2 and S5). From this, we conclude that there is an ordered structure in which the lateral dimensions of the molecules, the maximum and minimum feret diameters, increase proportionally, seemingly independent of the associated monomer units.

Biophysical studies of the fLuc mRNA multimer sample also showed a heterogeneous population, with MP detecting up to hexamer (6-mer) species and a mean population of ∼3.0 monomer units, corresponding to a trimer (3-mer). Relative to the monomer, the average volume and SBA detected by AFM were 3.7 and 3.1 times larger, respectively, with both showing a significant statistical difference, indicating substantial growth of the multimer species. Moreover, the average SBA is ∼3 times larger than that of the monomer sample, which aligns with the sample averaging to a trimer. The maximum feret diameter and mean height of the multimer sample resulted in a ∼1.6 and ∼1.1 increase from the monomer. Both of these increases are not statistically significant from the monomer, whilst the aspect ratio remains very similar at 0.797 (see Supplementary Table S3 and S7).

For both eGFP and fLuc mRNA, height increased significantly from monomer to dimer and multimer, but showed little or no further increase between dimer and multimer, indicating that initial dimer formation involves an increase in height, whereas subsequent association occurs predominantly through lateral expansion rather than vertical stacking. Consistent with this, SBA and volume differed significantly across all species and increased progressively from monomer to dimer to multimer, indicating an expanding immobilised footprint and continued accumulation of associated mRNA molecules, as supported by AFM images and mass photometry data. Although aspect ratio did not differ significantly between monomer and dimer, significant differences were observed between these species and multimers for fLuc mRNA, suggesting a transition from the more elongated morphology of monomers and dimers to a slightly more rounded multimer structure. eGFP and fLuc mRNA both followed similar trends across all parameters, with one notable difference in the maximum feret diameter between dimer and multimer populations (fLuc mRNA population had no significant difference, while eGFP mRNA population was significantly larger).

Through AFM and mass photometry analysis, we directly show that mRNA multimers adopt compact, ordered structures. mRNA multimers form similar laterally associated assemblies to the dimer and maintain the same overall shape as monomer units. There was no significant increase in the height of the whole structure immobilised on the mica surface. As observed during purification, the relative abundance of multimers of fLuc mRNA is lower than that of eGFP mRNA under the same conditions. Moreover, the extent of multimerisation (maximum detected multimer/n-mer) is greater for the eGFP mRNA, which forms up to a 9-mer in these conditions and has an average volume ∼9.0 times larger than the monomer, while fLuc mRNA forms up to a 6-mer and is only 3.7 times larger in volume. This indicates that for fLuc mRNA, self-association may be weaker under the same conditions as eGFP mRNA. This may be due to intramolecular folding of the secondary and tertiary mRNA structure suppressing quaternary structure formation (multimerisation), as intramolecular base-pairing sequesters regions that could form intermolecular bonds. Alternatively, the assembly of larger monomer units will occupy a larger footprint, leading to steric crowding/hindrance more readily, and therefore multimer growth will become self-limiting, resulting in the preferential formation of smaller assemblies.

## 4. Conclusions

In this study, we have developed a large-scale purification method using non-denaturing IP-RP chromatography for isolating discrete populations of mRNA monomers, dimers, and multimers. The use of 1 mM Mg^2+^ in the mobile phases stabilises intermolecular mRNA interactions, allowing separation of structures. Diafiltration of monomer, dimer and multimer samples using TFF was performed to remove Mg^2+^, acetonitrile and alkylamines. Following optimisation of the TFF, MP analysis and IP-RP HPLC showed that this did not dissociate or perturb the purified mRNA monomer, dimer, and multimer populations. We successfully used this methodology to produce purified mRNA monomer, dimer and mixed multimer populations for IVT-synthesised eGFP (931 nts) and fLuc mRNA (2121 nts). The ability to purify at scale will enable further downstream comparative biochemical and biophysical analysis of mRNA monomer, dimer and multimers, including their potential effects on protein expression, immunogenicity and encapsulation.

Further characterisation was performed using AFM to understand the structural and conformational makeup of each population of the mRNA higher-order structures. Automated analysis of the AFM images was performed to obtain volume, SBA, maximum feret diameter, height and aspect ratio measurements for mRNA monomer, dimer and multimer populations, providing further structural insight. AFM analysis revealed that the mRNA dimers and multimers formed compact, structured higher-order assemblies rather than end-to-end elongations. Volume, SBA, and maximum feret diameter increased with increasing higher-order structure and nucleotide length. The aspect ratio for all mRNA structures was between 0.7 and 0.8, indicating that mRNA species maintain a distinct shape, even during multimerisation. The volume of the mRNA dimers was more than double that of the monomers, consistent with the mRNA monomers adopting a more open conformation in the mRNA dimer. The separation of mRNA monomer from dimer and multimer species prior to AFM analysis enables analysis of discrete populations and therefore accurate morphometric measurement of conformation to be made. This improves quantitative comparison, and reduces analytical bias arising from overlapping size distributions. This work therefore provides a platform for the structural analysis of the quaternary structures of alternative mRNA constructs and their conformations under specific conditions.

## Data availability

The raw and processed atomic force microscopy (AFM) data supporting the findings of this study are openly available in the Atomic Force Microscopy Database (AFMDB), hosted within the Physical Sciences Data Infrastructure (PSDI) Community Data Collections - doi: 10.15131/shef.data.34995223. All other data supporting the conclusions of this article are included within the paper and its Supplementary Information files.

## Supplementary materials

Supplementary Data are available at *NAR* Online.

## CRediT authorship contribution statement

Alexandra L.J. Webb: Writing – review & editing, Writing – original draft, Visualisation, Validation, Methodology, Investigation, Formal analysis, Data curation, Conceptualisation. Harried Read: Writing – review & editing, Writing – original draft, Visualisation, Validation, Methodology, Investigation, Formal analysis, Data curation, Conceptualisation. Thomas C. Minshull: Writing – review & editing, Supervision, Data curation, Conceptualisation. Emma N. Welbourne: Writing – review & editing, Supervision, Conceptualisation. Prerna Bora: Resources, Investigation. Lily Brewster: Resources, Investigation. Manoj Pohare: Resources. Zoltan Kis: Writing – review & editing, Supervision. Sara Trabulo: Writing – review & editing, Supervision. Eivor Örnskov: Writing – review & editing, Supervision, Funding acquisition. Thomas E. Catley: Validation, Methodology, Investigation, Formal analysis, Writing – review & editing. Henrietta J. B. Smith: Resources, Methodology. Preksha Hooda: Investigation. Alice L. B. Pyne: Review & editing, Writing – original draft, Visualisation, Supervision, Funding acquisition, Conceptualisation. Mark J. Dickman: Writing – review & editing, Writing – original draft, Visualisation, Supervision, Funding acquisition, Conceptualisation.

## Acknowledgements

A.L.J.W is funded by an Engineering and Physical Sciences Research Council (EPSRC) Doctoral Training Partnership CASE Conversion award (EP/W524360/1) in collaboration with AstraZeneca. M.J.D and Z.K acknowledge funding from the Wellcome Leap R3 programme, Coalition for Epidemic Preparedness Innovations (CEPI), Biotechnology and Biological Sciences Research Council (BB/Y007514/1) and Innovate UK (Small Business Research Initiative, 10085632). MJD acknowledges additional funding from the Biotechnology and Biological Sciences Research Council (BB/Z515930/1). A.L.B.P. is funded by a UKRI Future Leaders Fellowship MR/W00738X/ and the Henry Royce Institute for Advanced Materials, funded through EPSRC grants EP/R00661X/1, EP/S019367/1, EP/P02470X/1 and EP/P025285/1. This work was supported through the UK-Southeast Asia Vaccine Manufacturing Research Hub. The Hub is funded by the Department of Health and Social Care using the UK Government’s International Development program (formerly “UK aid”) and is managed by the Engineering and Physical Sciences Research Council. The views expressed in this publication are those of the author(s) and not necessarily those of the Department of Health and Social Care. Graphical abstract created in BioRender. Dickman, M. (https://BioRender.com/uxmboll) is licensed under CC BY 4.0.

