## Supplementary Figures for "Purification and structural characterisation of mRNA monomers, dimers, and multimers using atomic force microscopy"

### Table of contents


| Supplementary information | Page number |
|---|---|
| Figure S1: Mass photometry calibration using High range Riboruler (Thermo Fisher Scientific) 500 – 6000 nucleotide RNAs. 1:150 dilution in 1X PBS. | 3 |
| Table S1: Mass photometry sizing of eGFP and fLuc mRNA fractions 1 (monomer), 2 (dimer) and 3 (multimers) with relative abundance (%). | 4 |
| Figure S2: Denaturing IP-RP HPLC analysis of eGFP monomer, dimer and multimer species. | 5 |
| Figure S3: Denaturing IP-RP HPLC analysis of fLuc monomer, dimer and multimer species. | 6 |
| Figure S4: AFM imaging of IVT-synthesised A) eGFP and B) fLuc mRNA with no prior non-denaturing, IP-RP or TFF purification. | 7 |
| Figure S5: AFM imaging of A) eGFP and B) fLuc monomer, dimer and multimer samples. | 8 |
| Figure S6: AFM measured heights of mRNA monomer, dimer and multimer samples, alongside post-hoc significance test results. | 9 |
| Table S2 AFM conformational parameters for eGFP mRNA monomer, dimer, multimer and unseparated populations. | 10 |
| Table S3 AFM conformational parameters for fLuc mRNA monomer, dimer, multimer and unseparated populations. | 11 |
| Table S4 Shapiro-Wilk Normality test to determine normality in eGFP mRNA AFM data n <5000., for $p < 0.05$ is not normal and $p > 0.05$ is normal. | 12 |
| Table S5 Post-hoc tests were performed to compare statistical significance between eGFP mRNA monomer, dimer and multimers | 13 |
| Table S6 Shapiro-Wilk Normality test was performed to determine normality in fLuc mRNA AFM data n <5000., for $p < 0.05$ is not normal and $p > 0.05$ is normal. | 14 |
| Table S7 Post-hoc tests were performed to compare statistical significance between eGFP mRNA monomer, dimer and multimers | 15 |

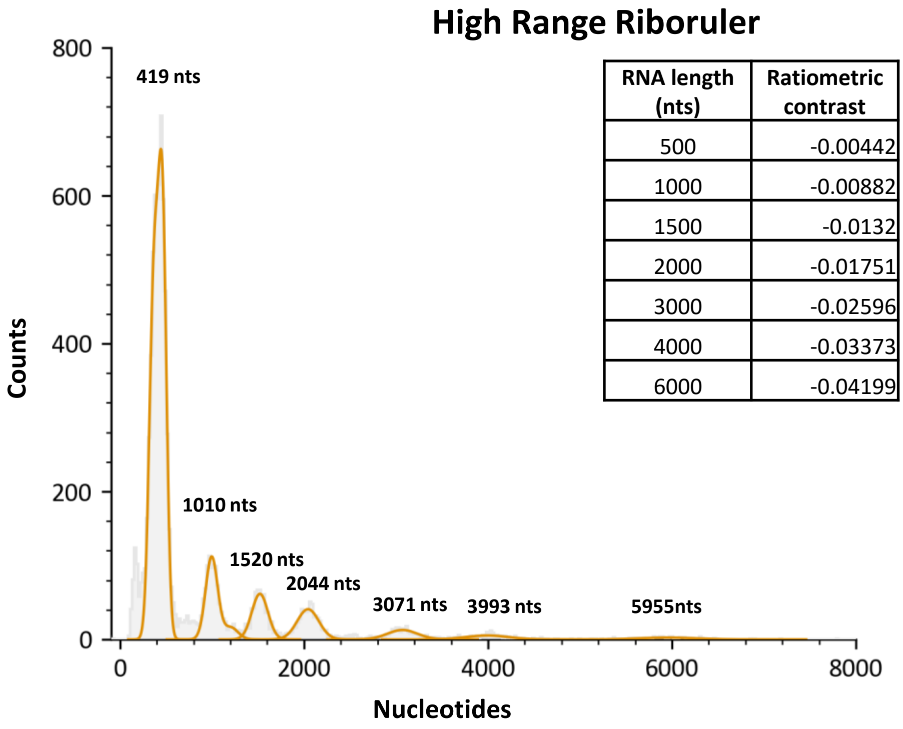


| RNA length (nts) | Ratiometric contrast |
|---|---|
| 500 | -0.00442 |
| 1000 | -0.00882 |
| 1500 | -0.0132 |
| 2000 | -0.01751 |
| 3000 | -0.02596 |
| 4000 | -0.03373 |
| 6000 | -0.04199 |

**Figure S1: Mass photometry calibration using High range Riboruler (Thermo Fisher Scientific) 500 – 6000 nucleotide RNAs. 1:150 dilution in 1X PBS.** Droplet dilution performed, and 10 µL of diluted RiboRuler loaded. Measurement taken over 60 seconds.

**Table S1: Mass photometry sizing of eGFP and fLuc mRNA fractions 1 (monomer), 2 (dimer) and 3 (multimers) with relative abundance (%).** eGFP samples were analysed pre-TFF and post-TFF with KR2i and µPulse systems. fLuc mRNA species was diafiltrated using the KR2i system only and then analysed.

| TFF system | | 1-mer | | 2-mer | | 3-mer | | 4-mer | | 5-mer | | 6-mer | | 7-mer | | 8-mer | | 9-mer | |
|---|---|---|---|---|---|---|---|---|---|---|---|---|---|---|---|---|---|---|---|
| | Fraction | % | nts | % | nts | % | nts | % | nts | % | nts | % | nts | % | nts | % | nts | % | nts |
| eGFP Pre-TFF | 1 | 100% | 913 | - | - | - | - | - | - | - | - | - | - | - | - | - | - | - | - |
| | 2 | 7.2% | 940 | 92.8% | 1861 | - | - | - | - | - | - | - | - | - | - | - | - | - | - |
| | 3 | 11.3% | 932 | 9.4% | 1927 | 16.9% | 2904 | 24.9% | 3806 | 14.7% | 4772 | 9.4% | 5627 | 6.1% | 6571 | 7.3% | 7873 | - | - |
| eGFP KR2i | 1 | 100% | 911 | - | - | - | - | - | - | - | - | - | - | - | - | - | - | - | - |
| | 2 | 10.4% | 954 | 89.6% | 1921 | - | - | - | - | - | - | - | - | - | - | - | - | - | - |
| | 3 | 12.8% | 890 | 8.8% | 1904 | 16.7% | 2873 | 22.9% | 3824 | 14.2% | 4797 | 7.7% | 5698 | 6.3% | 6560 | 5.9% | 7745 | 4.7% | 9532 |
| eGFP µPulse | 1 | 100% | 938 | - | - | - | - | - | - | - | - | - | - | - | - | - | - | - | - |
| | 2 | 36.1% | 936 | 63.9% | 1912 | - | - | - | - | - | - | - | - | - | - | - | - | - | - |
| | 3 | 30.1% | 924 | 24.7% | 1882 | 15.7% | 2868 | 9.6% | 3773 | 7.7% | 4629 | 6.1% | 5649 | 1.9% | 6567 | 2.2% | 7482 | 2.0% | 8603 |
| fLuc KR2i | 1 | 100% | 2050 | - | - | - | - | - | - | - | - | - | - | | | | | | |
| | 2 | 14.2% | 2028 | 77.2% | 4262 | 8.6% | 6430 | - | - | - | - | - | - | | | | | | |
| | 3 | 17.8% | 2055 | 21.6% | 4340 | 28.7% | 6549 | 18.0% | 8641 | 8.8% | 10714 | 5.4% | 12816 | | | | | | |

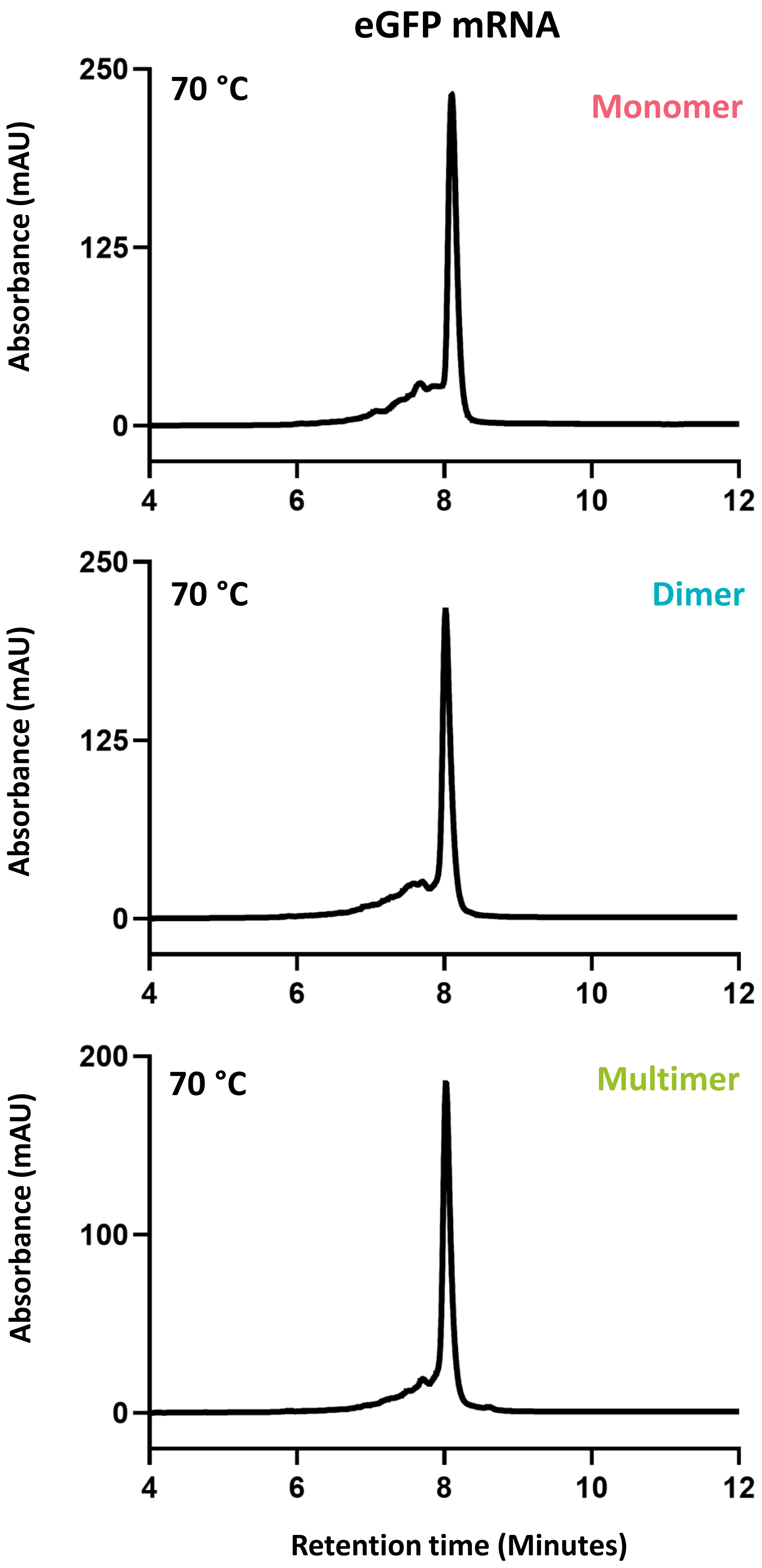


**Figure S2: Denaturing IP-RP HPLC analysis of eGFP monomer, dimer and multimer species.** IP-RP HPLC analysis was performed on U3000 and Vanquish HPLC systems (Thermo Fisher) using a DNAPac RP column (2.1 x 100 mm, Thermo Fisher). Mobile phase A was 0.1 M TEAA, and mobile phase B was 0.1 M TEAA, 25% ACN. The temperature was 70 °C. The gradient consisted of a linear step from 25-30% over 1 minute, a non-linear extension to 60% B over 14 minutes, and a 5-minute wash step at 100% B. Flow rate was 0.250 ml/min, and UV detection was at 260 nm.



Figure S2

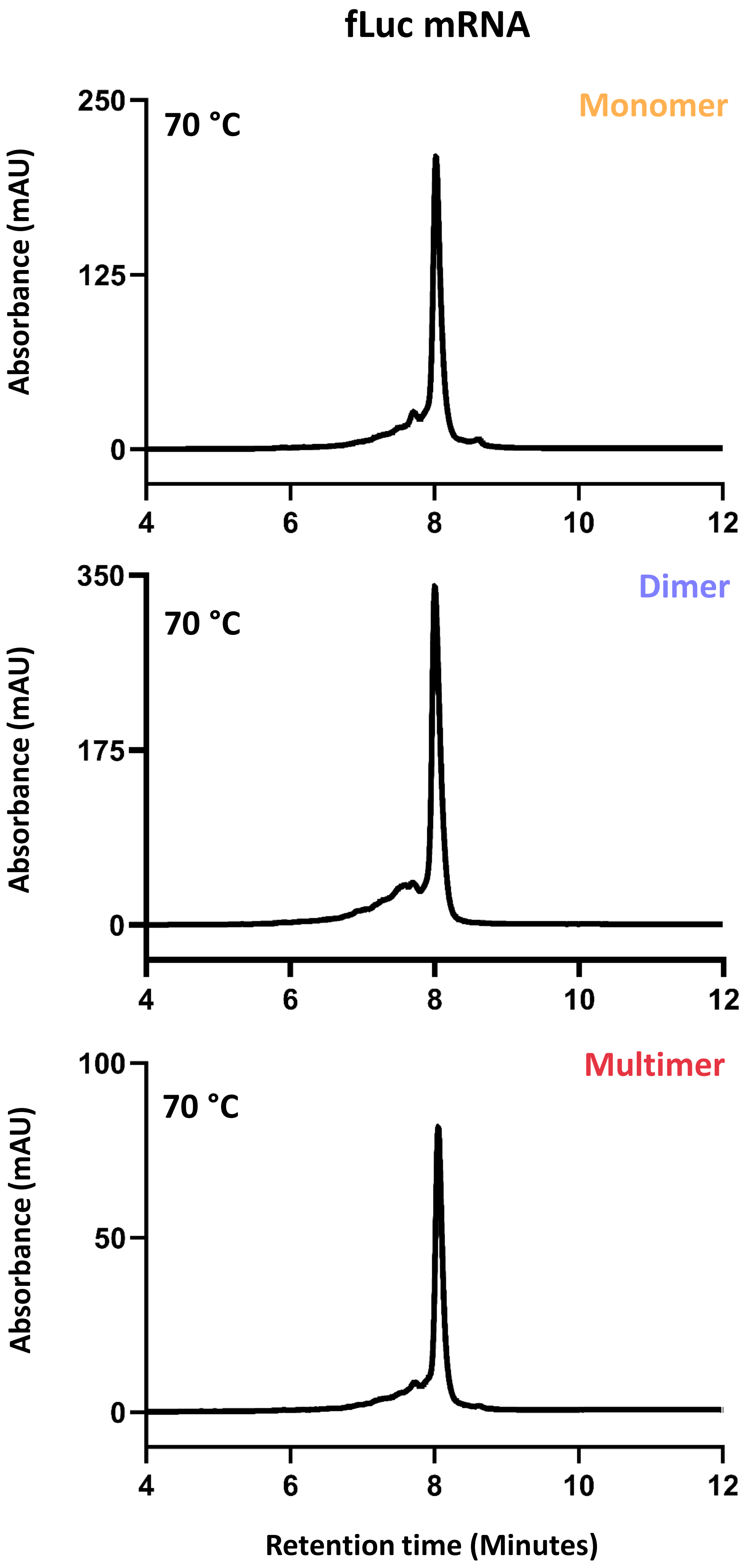


**Figure S3: Denaturing IP-RP HPLC analysis of fLuc monomer, dimer and multimer species.** IP-RP HPLC analysis was performed on U3000 and Vanquish HPLC systems (Thermo Fisher) using a DNAPac RP column (2.1 x 100 mm, Thermo Fisher). Mobile phase A was 0.1 M TEAA, and mobile phase B was 0.1 M TEAA, 25% ACN. The temperature was 70 °C. The gradient consisted of a linear step from 25-30% over 1 minute, a non-linear extension to 60% B over 14 minutes, and a 5-minute wash step at 100% B. Flow rate was 0.250 ml/min, and UV detection was at 260 nm.



Figure S3

A eGFP mRNA population

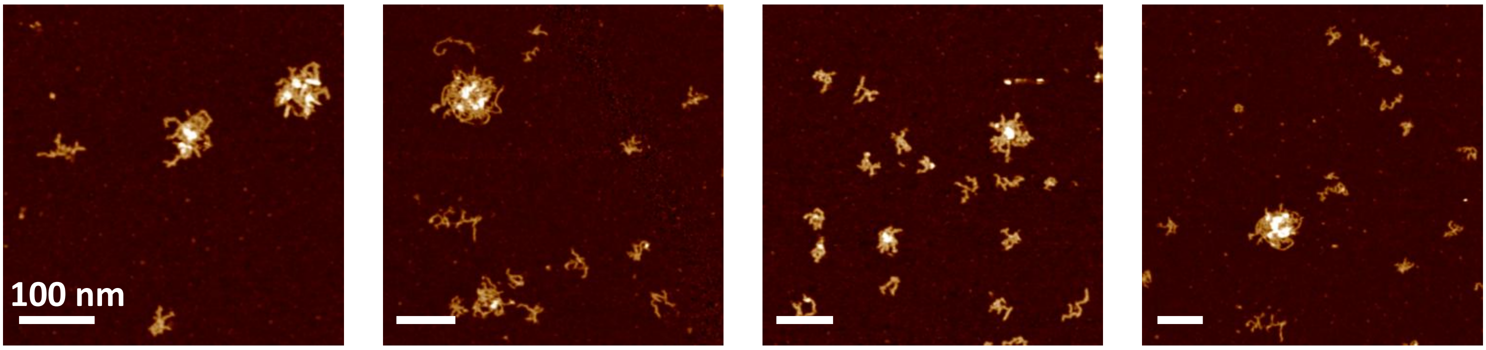


B fLuc mRNA population

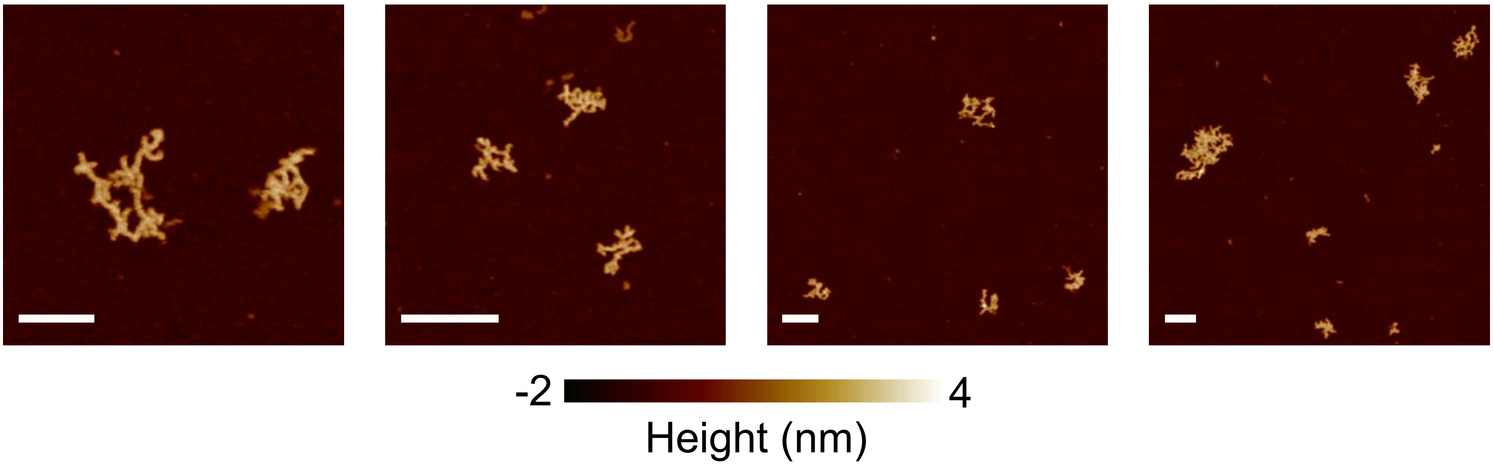


**Figure S4: AFM imaging of IVT-synthesised A) eGFP and B) fLuc mRNA with no prior non-denaturing, IP-RP or TFF purification.** Scale bar = 100nm.



Figure S4

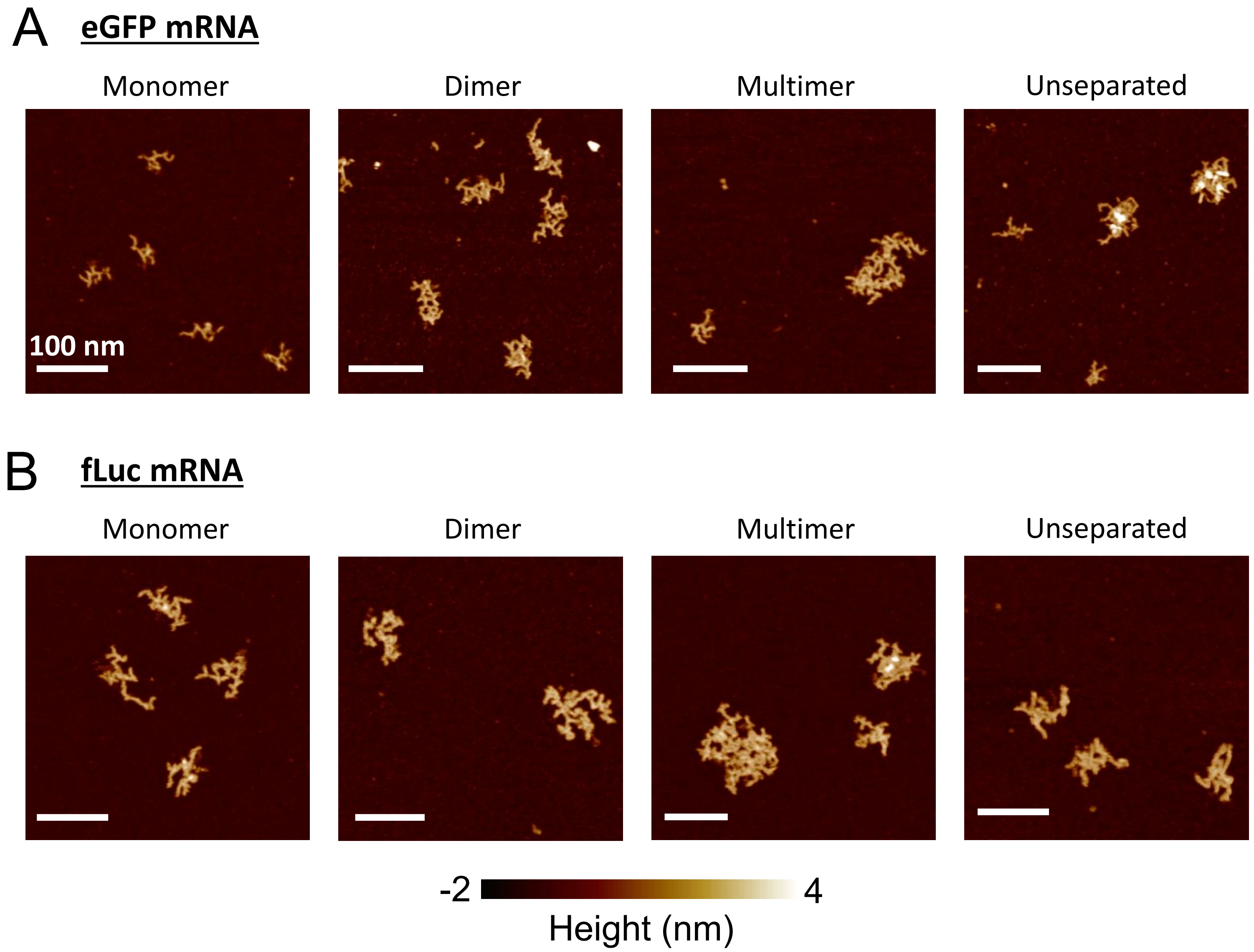


**Figure S5: AFM imaging of A) eGFP and B) fLuc monomer, dimer and multimer samples.** Scale bar = 100 nm for each AFM images, with all images being approximately similar sizes as to compare sizes of RNA species easily.



Figure S5

#### mRNA mean height

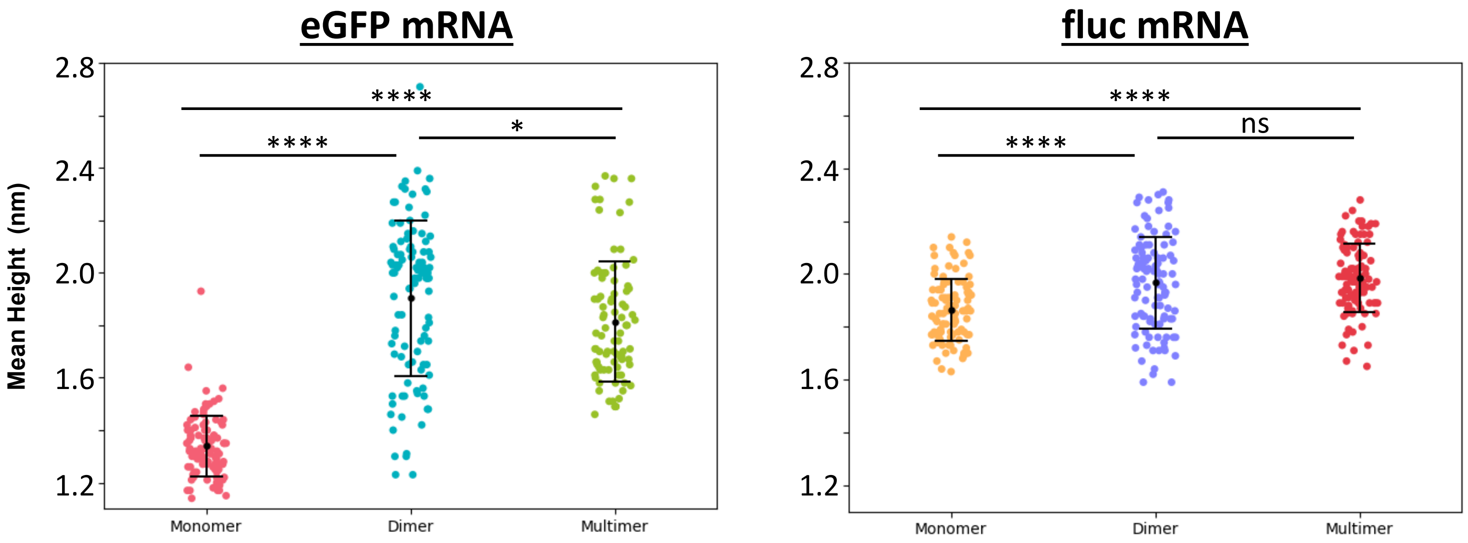


**Figure S6: AFM measured heights of mRNA monomer, dimer and multimer samples, alongside post-hoc significance test results.** Significance values can be found in tables S5 and S7.



Figure S6

**Table S2 AFM conformational parameters for eGFP mRNA monomer, dimer, multimer and unseparated populations.** Values are presented as mean ± SD, and are given to 3 significant figures.

| Sample | Volume ± SD ($nm^3$) | Smallest bounding area ± SD ($nm^2$) | Maximum Feret diameter ± SD (nm) | Aspect ratio ± SD |
|---|---|---|---|---|
| **Monomer** | 1220 ± 266 | 1690 ± 473 | 52 ± 8 | 0.735 ± 0.147 |
| **Dimer** | 3550 ± 1620 | 3280 ± 1270 | 72 ± 14 | 0.732 ± 0.157 |
| **Multimer** | 11000 ± 7500 | 10800 ± 7560 | 123 ± 42 | 0.753 ± 0.143 |
| **Unseparated** | 2241 ± 2580 | 2230 ± 1980 | 57 ± 21 | 0.745 ± 0.169 |

**Table S3 AFM conformational parameters for fLuc mRNA monomer, dimer, multimer and unseparated populations.** Values are presented as mean ± SD, and are given to 3 significant figures.

| Sample | Volume ± SD ($nm^3$) | Smallest bounding area ± SD ($nm^2$) | Maximum Feret diameter ± SD (nm) | Aspect ratio ± SD |
|---|---|---|---|---|
| **Monomer** | 4041 ± 1028 | 4156 ± 1368 | 80 ± 15 | 0.723 ± 0.142 |
| **Dimer** | 9959 ± 5601 | 9233 ± 4836 | 118 ± 30 | 0.723 ± 0.147 |
| **Multimer** | 14896 ± 12870 | 12750 ± 10856 | 130 ± 50 | 0.797 ± 0.148 |
| **Unseparated** | 8373 ± 14642 | 7897 ± 14461 | 98 ± 47 | 0.768 ± 0.146 |

**Table S4 Shapiro-Wilk Normality test to determine normality in eGFP mRNA AFM data, for $p < 0.05$ is not normal and $p > 0.05$ is normal.** Non-normally distributed data were analysed by the Kruskal-Wallis test, and normally distributed data were analysed by one-way ANOVA. All values are given to 3 significant figures.

### eGFP mRNA

| Parameter | Monomer | Dimer | Multimer |
|---|---|---|---|
| Smallest Bounding Area | $p = 2.02 \times 10^{-6}$ | $p = 6.86 \times 10^{-14}$ | $p = 2.32 \times 10^{-8}$ |
| Volume | $p = 4.80 \times 10^{-11}$ | $p = 6.00 \times 10^{-11}$ | $p = 2.61 \times 10^{-8}$ |
| Max Feret | $p = 0.134$ | $p = 3.06 \times 10^{-7}$ | $p = 5.96 \times 10^{-4}$ |
| Aspect Ratio | $p = 4.25 \times 10^{-4}$ | $p = 4.89 \times 10^{-2}$ | $p = 9.04 \times 10^{-2}$ |
| Mean Height | $p = 6.65 \times 10^{-6}$ | $p = 8.26 \times 10^{-3}$ | $p = 2.80 \times 10^{-4}$ |

**Table S5. Post-hoc tests were performed to compare statistical significance between eGFP mRNA monomer, dimer and multimers**. Significance was assigned as such, $p < 0.0001$ ****, $p < 0.001$ ***, $P < 0.01$ **, $P < 0.05$ *. All parameters showed statistically significant differences to a varying degree, apart from the aspect ratio, which remained non-significant between the monomer, dimer and multimer samples. All values are given to 3 significant figures.

### eGFP mRNA

| Parameter | Monomer vs Dimer | Monomer vs Multimer | Dimer vs Multimer |
|---|---|---|---|
| **Height** | 0 → 0 **** | 0 → 0 **** | $1.95 \times 10^{-2}$ * |
| **Smallest bounding area** | $2.01 \times 10^{-2}$ * | 0 → 0 **** | 0 → 0 **** |
| **Volume** | $2.90 \times 10^{-4}$ *** | 0 → 0 **** | 0 → 0 **** |
| **Maximum Feret diameter** | $1.69 \times 10^{-8}$ **** | 0 → 0 **** | 0 → 0 **** |
| **Aspect ratio** | 0.986 | 0.714 | 0.613 |

**Table S6 Shapiro-Wilk Normality test was performed to determine normality in fLuc mRNA AFM data for $p < 0.05$ is not normal and $p > 0.05$ is normal.** Non-normally distributed data were analysed by the Kruskal-Wallis test, and normally distributed data were analysed by one-way ANOVA.

### fLuc mRNA

| | Monomer | Dimer | Multimer |
|---|---|---|---|
| **Smallest Bounding Area** | $p = 5.27 \times 10^{-9}$ | $p = 9.16 \times 10^{-10}$ | $p = 2.56 \times 10^{-11}$ |
| **Volume** | $p = 3.05 \times 10^{-14}$ | $p = 3.85 \times 10^{-12}$ | $p = 8.40 \times 10^{-12}$ |
| **Max Feret** | $p = 1.34 \times 10^{-5}$ | $p = 4.09 \times 10^{-6}$ | $p = 1.02 \times 10^{-6}$ |
| **Aspect Ratio** | $p = 0.271$ | $p = 0.147$ | $p = 5.84 \times 10^{-5}$ |
| **Mean Height** | $p = 0.124$ | $p = 0.147$ | $p = 0.398$ |

**Table S7 Post-hoc tests were performed to compare statistical significance between eGFP mRNA monomer, dimer and multimers**. Significance was assigned as such, $p < 0.0001$ ****, $p < 0.001$ ***, $P < 0.01$ **, $P < 0.05$ *. All parameters showed statistically significant differences to a varying degree, apart from the aspect ratio, which remained non-significant between the monomer, dimer and multimer samples. All values are given to 3 significant figures.

### fLuc mRNA

| Parameter | Monomer vs Dimer | Monomer vs Multimer | Dimer vs Multimer |
|---|---|---|---|
| **Height** | $1.33 \times 10^{-6}$ **** | < 0.0001 **** | 0.6070 |
| **Smallest bounding area** | $2.03 \times 10^{-7}$ **** | < 0.0001 **** | $7.93 \times 10^{-4}$ *** |
| **Volume** | $6.25 \times 10^{-7}$ **** | < 0.0001 **** | $4.33 \times 10^{-5}$ **** |
| **Maximum Feret diameter** | $1.77 \times 10^{-13}$ **** | < 0.0001 **** | $6.13 \times 10^{-2}$ |
| **Aspect ratio** | 0.979 | $8.38 \times 10^{-4}$ *** | $1.33 \times 10^{-3}$ ** |
